# An endoplasmic reticulum resident molecular chaperone, GRP170, prevents stress-induced glomerular injury

**DOI:** 10.64898/2026.01.28.702322

**Authors:** Aidan Porter, Hannah E. Vorndran, Allison Marciszyn, Anuradha Iyer, Thomas R. Kleyman, Edward A. Fisher, Roderick J. Tan, Jeffrey L. Brodsky, Teresa M. Buck

## Abstract

The glomerulus, a unique capillary network in the nephron, filters an entire blood volume approximately 300 times a day. Specialized epithelial cells known as podocytes form a critical component of the glomerular filtration barrier, and diseases linked to podocyte injury include minimal change disease, focal segmental glomerulosclerosis, and diabetic kidney disease. Because podocytes are terminally differentiated, their ability to respond to external stress is critical. The unfolded protein response (UPR), a cellular stress pathway, is associated with glomerular injury, although the role of the UPR in glomerular injury is undefined. The UPR is initially protective, leading to upregulation of molecular chaperones, a class of proteins that promote protein folding and are required to survive oxidative and ischemic injury. An unresolved UPR, however, leads to apoptosis. We previously found that one molecular chaperone, GRP170, provides protection against acute kidney injury since GRP170 depletion led to UPR induction and widespread kidney injury. Here we generated a new podocyte specific GRP170 knock out mouse (GRP170^Pd-/-^). Surprisingly, GRP170^Pd-/-^ mice were born healthy, and podocyte development appeared normal. Within a month, however, the knockout mice exhibited profound glomerular injury manifesting as proteinuria, hypoalbuminemia, hyperlipidemia, and kidney injury. Concomitant with glomerular injury, we observed increased expression of the pro-apoptotic UPR target, CHOP, in podocytes. Together, our new model not only defines a protective role for GRP170 against glomerular injury but also provides a new model to test the therapeutic potential of small molecule UPR modulators to treat glomerular injury.

## INTRODUCTION

The nephron is the fundamental structural and functional unit of the kidney and is responsible for fluid and electrolyte balance, waste excretion, blood pressure regulation, and mineral and amino acid metabolism. The human kidney contains approximately one million nephrons, which collectively filter up to 180 liters of plasma daily. Filtration occurs in the glomerulus, a specialized structure that permits water and small solutes to enter the urinary space, while retaining most plasma proteins and cellular macromolecules in the circulation (Scott and Quaggin 2015; Pollak et al. 2014). Selectivity is conferred by the glomerular filtration barrier—a dynamic three-layered structure composed of fenestrated endothelial cells and glomerular epithelial cells, which are known as podocytes. These cell types are separated from one another by an extracellular matrix, known as the glomerular basement membrane (GBM). Slit diaphragms, specialized membrane-like junctions between interdigitating podocyte foot processes, work in concert with the GBM to optimize filtration.

The loss of glomerular integrity is catastrophic, leading to hematuria, proteinuria, and progressive glomerular and renal tubular disease. While endothelial cells and the GBM are also susceptible to damage, mounting evidence indicates that podocyte injury is a key pathogenic driver in many glomerular diseases, including minimal change disease, focal segmental glomerulosclerosis (FSGS), and diabetic kidney disease (DKD) (Lv et al. 2025; Cybulsky et al. 2011; Benzing and Salant 2021). Like other cell types that require a diverse repertoire of critical secreted and transmembrane proteins, podocytes are particularly reliant on endoplasmic reticulum (ER) function, since this is the site of membrane and secreted protein synthesis, folding, and maturation(Ren et al. 2018; Yoshida et al. 2021; Cybulsky 2017; Cheng et al. 2017). In addition, podocytes exhibit limited regenerative capacity, which makes them especially vulnerable to injury (Hartleben et al. 2010; Taniguchi and Yoshida 2015; Yang et al. 2023).

When confronted with injury, the podocyte—along with most other cell types—engages a cell stress pathway known as the Unfolded Protein Response (UPR) (Gonçalves et al. 2018; Hu et al. 2023; Navarro-Betancourt and Cybulsky 2022; Inagi et al. 2005). The UPR is triggered by the accumulation of misfolded proteins, calcium dysregulation, oxidative stress, altered protein glycosylation, nutrient deprivation, and lipid imbalance (Kettel and Karagöz 2024; Hetz et al. 2020). Three ER-membrane receptors—IRE1, ATF6, and PERK—detect ER stress and initiate distinct yet coordinated downstream signaling cascades to restore protein homeostasis (proteostasis), or, if stress persists, promote cell death. IRE1 and ATF6 function together to drive early protective responses, upregulating the production of ER molecular chaperones, which facilitate protein folding, expanding ER capacity, which reduces the concentration of misfolded proteins in the ER, and reducing global protein synthesis, so the ER protein folding machinery is not overburdened (Hetz et al. 2020; Porter, Brodsky, et al. 2022). While the PERK pathway is initially protective(Liu et al. 2024), sustained PERK activation under severe or unrelenting stress selectively induces the expression of the proapoptotic transcription factors ATF4 and CHOP (Iurlaro and Muñoz-Pinedo 2016; Kettel and Karagöz 2024).

In contrast to the characterized ER molecular chaperones that maintain ER proteostasis, the role of another abundant and conserved chaperone, GRP170, is less defined. GRP170 (also known as ORP150 and HYOU1) acts as a nucleotide exchange factor (NEF) for the ATP-requiring ER lumenal Hsp70, BiP (Tyson and Stirling 2000). GRP170 also exhibits “holdase” function, retaining non-native, aggregation-prone proteins in a soluble state (Easton et al. 2000; Chen et al. 1996; Behnke et al. 2016; Craven et al. 1996). Although only a handful of GRP170 substrates have been identified, we established that GRP170 is required for the degradation of unassembled subunits of the Epithelial Sodium Channel (ENaC), a heterotrimer that maintains salt and water balance in the kidney by re-absorbing sodium in the distal nephron (Buck et al. 2013; Sukhoplyasova et al. 2023; Buck et al. 2017).

To more generally understand how GRP170 contributes to kidney physiology, we previously generated a doxycycline-inducible, nephron-tubule-specific, GRP170 KO mouse (GRP170^NT-/-^) (Porter, Nguyen, et al. 2022; Porter et al. 2025). GRP170 deletion resulted in hyperkalemia, hyponatremia, and an acute kidney injury (AKI)-like phenotype that was first accompanied by induction of the protective arms of the UPR but ultimately characterized by PERK- and caspase-dependent renal cell death; in fact, multiple lines of evidence link the UPR to AKI (Porter, Brodsky, et al. 2022). To differentiate whether renal cell death arose from altered salt/water homeostasis or UPR induction, we maintained electrolyte levels by placing GRP170^NT-/-^ mice on a high sodium diet. While several injury phenotypes were improved, the UPR was unaffected by the high sodium diet, suggesting that full correction of AKI-like symptoms will ultimately also require the administration of UPR modulators (Porter et al. 2025). Indeed, when GRP170 was depleted over time in mouse embryonic fibroblasts (MEFs)—and consistent with the importance of GRP170 as a BiP regulator—misfolded proteins began to aggregate and latent pools of BiP were activated as GRP170 levels fell. Nevertheless, cell death was ultimately triggered by CHOP/caspase-dependent cell death (Mann et al. 2024).

Beyond AKI, the UPR has also been implicated in glomerular injury, including minimal change disease, membranous nephropathy, FSGS, and diabetic nephropathy (Yang et al. 2023; Tao et al. 2016; Markan et al. 2009; Madhusudhan et al. 2015; Chung et al. 2023). In parallel, GRP170 has emerged as one of the ER molecular chaperones that is amplified in proteinuric kidney disease (Wang et al. 2019; Inagi et al. 2005; 2008; Lindenmeyer et al. 2008). Yet, whether GRP170 is directly required to maintain podocyte proteostasis has not been reported. To this end, we developed a new podocyte-specific GRP170 KO mouse (GRP170^Pd-/-^). To our surprise, GRP170^Pd-/-^ mice are viable and exhibit a mild phenotype at post-natal day 13 (P13), suggesting that podocyte development is largely preserved in the absence of GRP170. In contrast, by P28, we observed a rapid decline in podocyte function, manifesting as glomerular filtration barrier defects and subsequent proteinuria, hypoalbuminemia, hyperlipidemia, and kidney injury.

## RESULTS

### GRP170 is required to maintain podocyte function

To more definitively link ER stress and the UPR to glomerular injury—and given the importance of GRP170 in maintaining ER homeostasis in other animal models (Ye et al. 2013; Kitano et al. 2004; Miyazaki et al. 2002; Zhang et al. 2025)—we generated a GRP170-deficient podocyte-specific mouse model. A mouse harboring a floxed allele at the GRP170-encoding locus was crossed to the *NPHS2-*Cre mouse, which expresses Cre recombinase exclusively in podocytes(Moeller et al. 2003; Porter, Nguyen, et al. 2022). The resulting GRP170^Pd-/-^ mice were viable, obtained at the expected Mendelian ratios, and were phenotypically indistinguishable from littermate controls at birth. The GRP170^Pd-/-^ and heterozygous GRP170^Pd+/-^ animals were also indistinguishable from GRP170^Pd+/+^ animals at P13 (see e.g. Figure 1A). Moreover, serum BUN and creatinine concentrations were similar across genotypes (Figure 1B), and plasma electrolytes, acid-base balance, and hemoglobin were also comparable at this stage (Figure 1C). Proteinuria, which is another direct and often early manifestation of podocyte injury and can, itself, accelerate the progression of kidney disease (Benzing and Salant 2021; Kopp et al. 2020), as well as hypoalbuminemia, were also absent (Figure 1D). In contrast to preserved glomerular function in GRP170^Pd-/-^ mice, germline deletion other cytoprotective genes, such as *Tac1* and *Cdc42*, compromise podocyte development, leading to congenital nephrotic syndrome and perinatal mortality within days (Kim et al. 2014; Scott et al. 2012).

**Figure 1:**
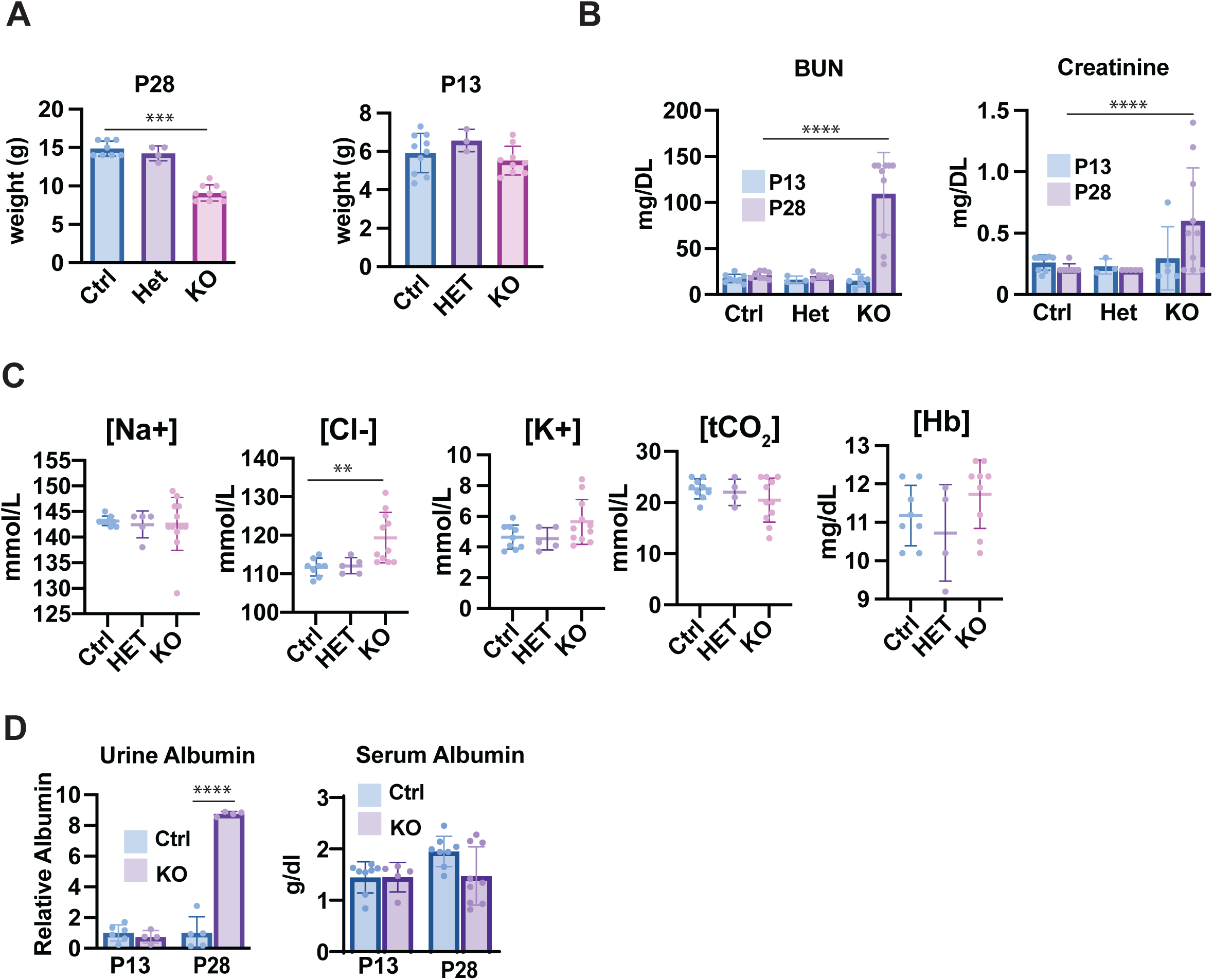
GRP170^PD-/-^ mice have evidence of podocyte and nephron injury by P28. GRP170^PD-/-^ and control P13 and P28 mice were weighed (**A**) and blood chemistry (**B, C**), and serum and urine albumin (**D**) were measured. Data represent the means +/- SD; n=5-10. **P < 0.01, ***P < 0.001, ****P < 0.0001. Statistical significance determined by 1-way ANOVA followed by Tukey’s multiple-comparison test.

Phenotypes associated with most renal diseases, including those that affect podocyte function, worsen over time, and in other podocyte-injury mouse models select phenotypes followed this trend (Scott et al. 2012; Roselli et al. 2004; Putaala et al. 2001). Therefore, we examined phenotypes in each cohort of animals at P28. At this age, the GRP170^Pd-/-^ animals weighed significantly less than their littermates and had significantly higher serum levels of standard AKI markers, i.e., BUN and creatinine, than GRP170^Pd+/+^ and GRP170^Pd+/-^ animals (Figure 1A, B). Plasma analyses also revealed a trend toward lower albumin and bicarbonate and higher potassium and hemoglobin levels in GRP170^Pd-/-^ mice than in their littermates (Figure 1C, D). In contrast, plasma chloride was substantially elevated in the knockout (KO) animals, likely reflecting a hypoalbuminemia-driven compensatory response to maintain plasma electroneutrality (Seifter 2014). Consistent with podocyte injury, urine albumin was also significantly elevated by P28 (Figure 1D). These findings are consistent with intravascular volume contraction due to urinary protein loss, resulting in hemoconcentration, weight loss, and impaired tubular function in the absence of GRP170. Collectively, these data indicate that GRP170 is critical for postnatal glomerular maintenance.

### Podocyte-specific loss of GRP170, and glomerular injury, is accompanied by increased serum cholesterol and apolipoprotein B

Because hyperlipidemia and lipiduria are defining features of nephrotic syndrome, we also measured plasma lipids in GRP170^Pd-/-^ and control mice at P13 and P28 (Figure 2A) (Ponticelli and Moroni 2023; Zaiou et al. 1998a). As anticipated, triglyceride (TG) and cholesterol levels were comparable in P13 mice regardless of genotype but were significantly elevated in P28 GRP170^Pd-/-^ mice. Because serum cholesterol and TG are associated with hepatic and intestinal-synthesized apolipoprotein B (ApoB), we also quantified plasma levels of this lipoprotein. More specifically, we blotted for both the ApoB100 and ApoB48 isoforms, which serve as the core structural proteins of very low-density lipoproteins/low density lipoproteins (VLDL/LDL) and chylomicrons, respectively (Brodsky and Fisher 2008). Consistent with the TG and cholesterol measurements, ApoB100 and ApoB48 levels were comparable amongst the two genotypes at P13 but dramatically increased in P28 GRP170^Pd-/-^ mice relative to GRP170^Pd+/+^ animals (Figure 2B).

**Figure 2:**
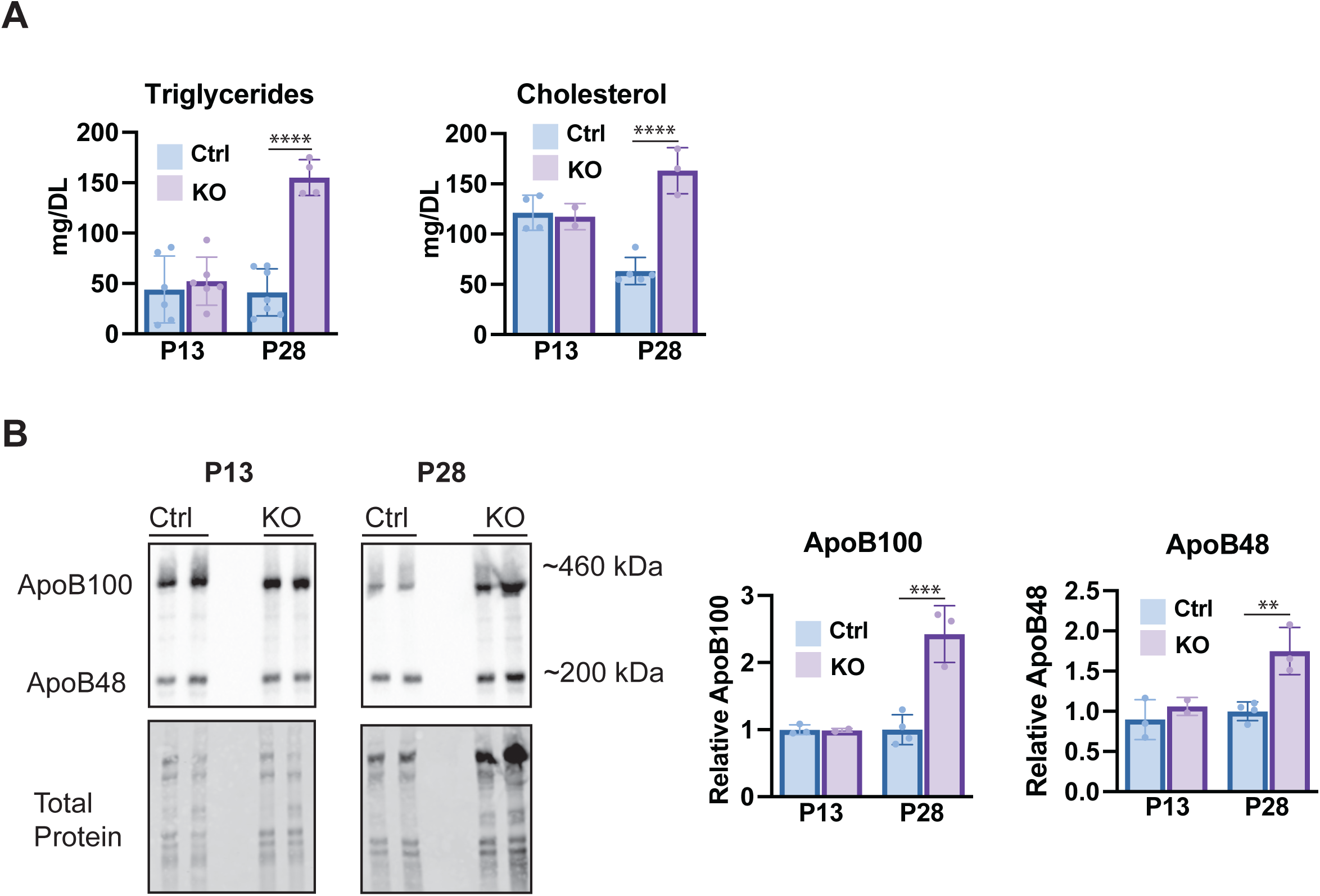
Loss of GRP170 results in lipid dysregulation in the GRP170^Pd-/-^ mice. (A) Plasma triglycerides and cholesterol levels from P13 or P28 GRP170^Pd-/-^ mice were determined as described in *Materials and Methods*. (B) Immunoblotting ApoB from P13 or P28 GRP170^PD-/-^ and control mouse plasma. Data represent the means +/- SD; n=3-6. **P < 0.01, ***P < 0.001, ***P < 0.0001. Statistical significance determined by 1-way ANOVA followed by Tukey’s multiple-comparison test.

### Renal histology of GRP170^Pd-/-^ mice is consistent with podocyte injury

To better assess the extent of glomerular and tubular injury in GRP170^Pd-/-^ mice, fixed kidney sections from P13 and P28 GRP170^Pd-/-^ mice and controls were subject to hematoxylin and eosin (H&E) staining. As shown in Figure 3, renal histology in control GRP170^Pd-/-^ mice at P13 appeared normal. In contrast, the kidneys of P28 GRP170^Pd-/-^ mice selectively displayed markedly distorted podocyte and tubular architecture. Many glomeruli were small and collapsed with obliterated capillary loops and evidence suggestive of early sclerosis. Compared to healthy tissue, the renal tubules were also dilated, fewer in number, and more widely spaced with increased interstitial cellularity. The tubular lumens additionally contained numerous proteinaceous casts (marked with an * in Figure 3, bottom right panel). These findings, which resemble those observed in progressive proteinuric glomerular diseases, provide a structural basis for the severe proteinuria arising from podocyte dysfunction in the absence of GRP170 (Hill et al. 2001; Hebert et al. 2000; Antonovych, T. T., Mostofi, F. K. 1980).

**Figure 3.**
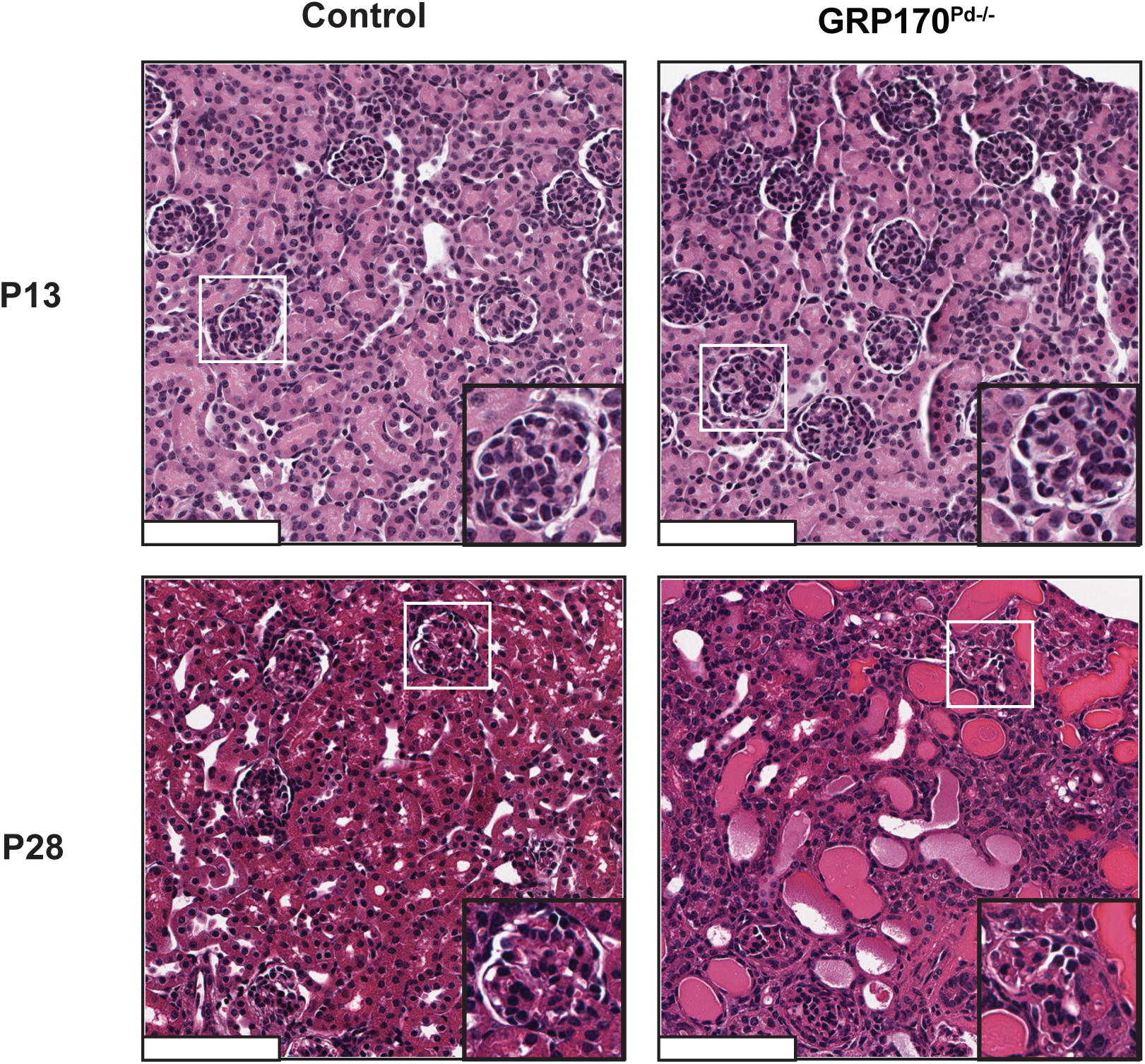
Renal histology of GRP170^Pd-/-^ mice confirms podocyte and renal tubular injury. Hematoxylin and eosin (H&E)-stained kidney sections from P13 and P28 GRP170^Pd-/-^ mice and controls were examined. Scale bars: 200 µm.

### GRP170 deletion causes progressive podocyte injury and slit diaphragm degeneration

To more precisely visualize how GRP170 deletion compromises podocyte and glomerular structure, we performed indirect immunofluorescence microscopy and examined WT1 (Wilms’ tumor gene 1), a transcription factor critical to podocyte development, differentiation, and maintenance (Dong et al. 2015). Because WT1 is expressed in mature podocytes, the number of glomerular WT1 positive cells is both a reliable surrogate for podocyte number and integrity as well as a sensitive indicator of podocyte injury. As anticipated based on the data shown in Figure 3, P13 GRP170^Pd-/-^ and control mice displayed similar numbers of WT1-positive cells, whereas P28 GRP170-deficient mice had ∼1/3 fewer WT1-positive cells than controls (Figure 4A, B). A marked reduction in the number of WT1-positive cells is consistent with the small, collapsed glomeruli seen in the H&E staining shown above.

**Figure 4:**
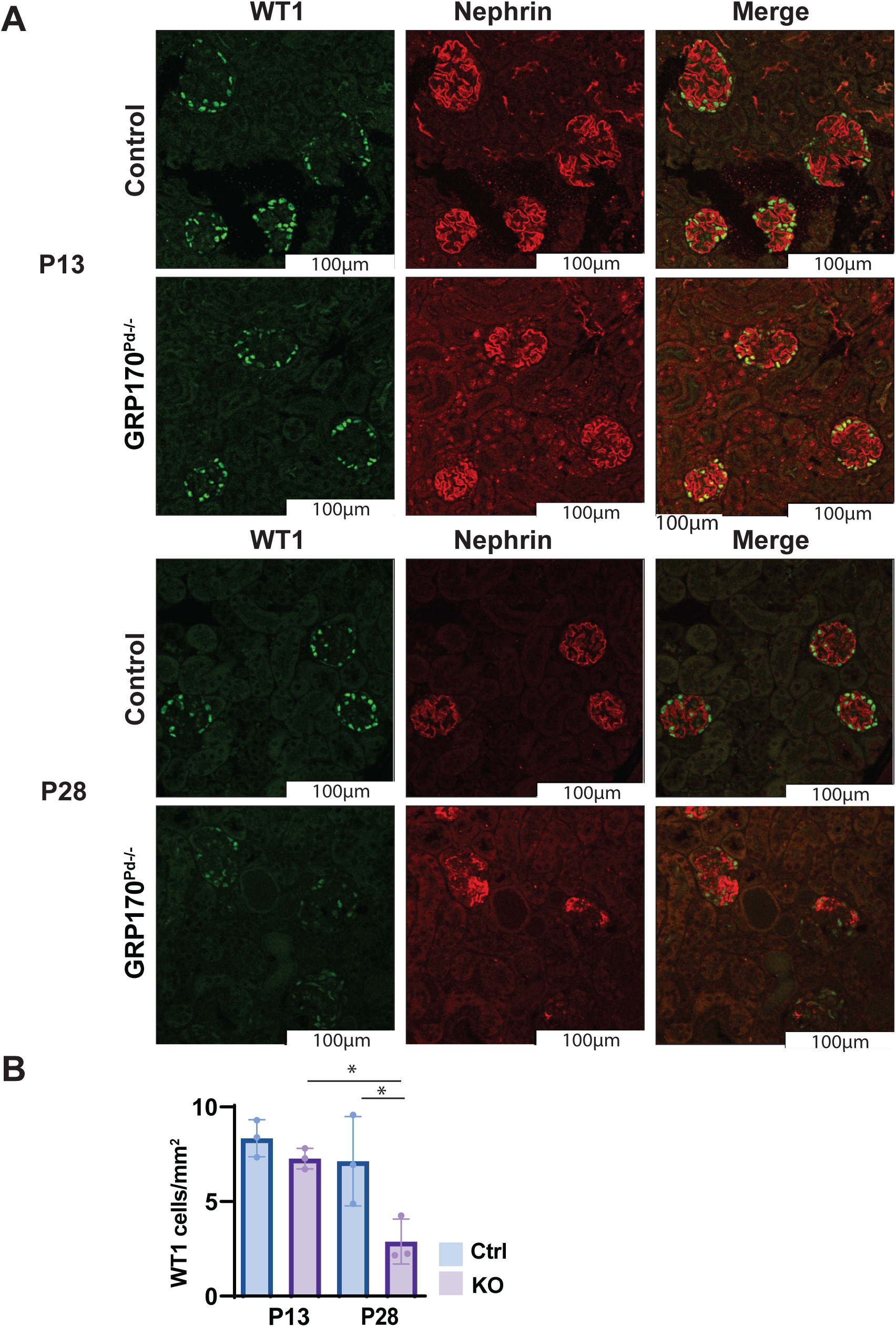
Loss of GRP170 causes deterioration of the podocyte slit diaphragm and rarefaction of podocytes within the glomerulus. (**A**) Representative immunofluorescence images depicting WT1 (podocyte nuclear marker) and nephrin (slit diaphragm). Scale: 100 µm. (**B**) Quantification of WT1-positive cells per glomerulus. Data are presented as mean ± SD (n = 3 mice per genotype at P13 and P28). For each kidney section, WT1-positive cells were counted in 3–9 glomeruli. *P < 0.05.

To more precisely assess filtration barrier integrity, we also performed immunofluorescence staining for nephrin (Figure 4A). As a transmembrane protein that spans adjacent podocyte foot processes to form the slit diaphragm, nephrin is a critical component of the GBM and assists in regulating the podocyte actin cytoskeleton and maintaining cell polarity (Williamson et al. 2025). Mutations in *NPHS1*, which encodes nephrin, cause congenital nephrotic syndrome, and diminished nephrin expression is a common feature in proteinuric kidney diseases (L. Liu et al. 2001; Verma et al. 2018; Benzing and Salant 2021). In healthy glomeruli with intact GBM architecture, nephrin staining forms a continuous linear pattern along the glomerular capillary loop, and indeed normal nephrin staining was obtained in sections from P13 GRP170^Pd-/-^ mice. By P28, however, nephrin staining was significantly depleted in podocytes in GRP170^Pd-/-^ mice. Since we observed fewer podocytes in P28 GRP170^Pd-^ mice and given the distorted glomerular architecture and slit diaphragm integrity, we also performed immunofluorescence against endomucin and PDGFRβ, which mark endothelial capillary and mesangial cells, respectively (C. Liu et al. 2001; Samulowitz et al. 2002; Seifert et al. 1998). In each case, the kidneys of GRP170-deficient and control animals were indistinguishable at P13. By P28, however, endomucin and PDGFβ signal were also less prominent in GRP170^Pd-/-^ mice (Supplemental Figure S1A, B), consistent with indiscriminate glomerular cell death. In aggregate, the staining patterns we observed suggest that the glomeruli and slit diaphragm undergo progressive degeneration when podocyte GRP170 is absent, and downstream consequences of this effect are seen in other renal cell types.

### Podocyte ultrastructure reveals early signs of podocyte injury in P13 GRP170^Pd-/-^ mice

The apparent health, absence of proteinuria, and normal renal morphology observed using light and confocal microscopy suggest that podocyte development and function are intact in the P13 GRP170^Pd-/-^ mouse. In its earliest stages, however, podocyte injury, as in minimal change disease, can be indiscernible by traditional microscopy techniques (Yu et al. 2018). Therefore, to determine whether subtle abnormalities are already present in P13 animals, and to better define the onset of podocyte dysfunction in the absence of GRP170, we examined kidney sections from P13 GRP170^Pd-/-^ and control mice by SEM (scanning electron microscopy) and TEM (transmission electron microscopy). As shown in Figure 5, we found that GRP170-deficient mice displayed thickening (see blue boxes) and mild focal effacement of their foot processes (see red box), indicating that GRP170 function supports podocyte integrity even at this early stage. Because the GRP170^Pd-/-^ mouse is a constitutive knockout, it is unclear whether podocyte injury arises prenatally or postnatally. However, the absence of proteinuria at P13 suggests that podocyte damage is subtle.

**Figure 5:**
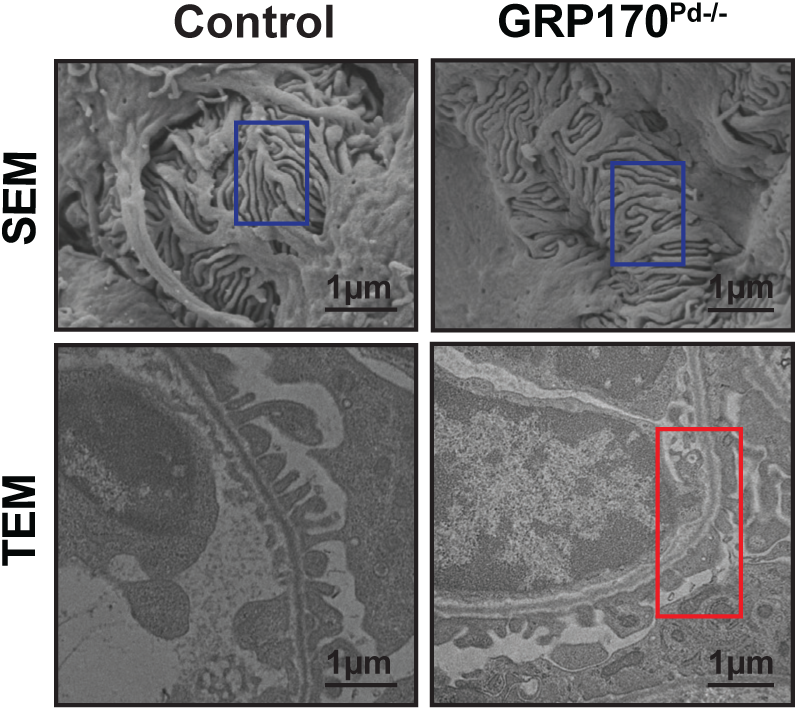
GRP170 deletion causes ultrastructural podocyte degeneration in P13 GRP170^Pd-/-^ mice. Representative SEM and TEM of kidney sections from P13 GRP170^Pd-/-^ mice showed thickening (blue boxes) and focal effacement (red box) of interdigitating foot processes.

### Progressive UPR induction in the GRP170^Pd-/-^ mouse

As highlighted previously(Porter, Nguyen, et al. 2022; Porter et al. 2025; Mann et al. 2024), the loss of Grp170 is accompanied by UPR induction. To provide direct evidence of the UPR in GRP170-deficient podocytes, we stained kidney sections for BiP, a proximal target of the IRE1 arm of the UPR, and CHOP, a downstream pro-apoptotic factor induced by the PERK branch of this pathway (Ron and Walter 2007). We also co-stained for synaptopodin, a protein localized to podocyte foot processes that is critical for cytoskeletal dynamics along with the integrity of the glomerular filtration barrier (Yu et al. 2018; Ning et al. 2021). As anticipated, synaptopodin appeared in a ring-like staining pattern, which demarcated the podocyte boundary. At P13, BiP staining was more prominent in these podocytes in sections from GRP170^Pd-/-^ animals compared to the controls, consistent with early induction of the protective UPR (Figure 6A). In contrast, CHOP staining was largely absent at P13, suggesting the destructive branch of the UPR remained silent at P13, but by P28 BiP levels returned to near control levels in GRP170^Pd-/-^ mice, consistent with the transient activation of BiP we observed in MEFs (Mann et al. 2024). In contrast, CHOP staining was pronounced at P28 in the knockout mice. This outcome likely reflects progression of podocyte injury, a sustained UPR, and initiation of the apoptotic pathway. Notably, CHOP expression appeared largely confined to the glomeruli, even though the tubules of P28 GRP170^Pd-/-^ animals exhibited marked structural damage on H&E staining and widespread apoptosis, as indicated by the abundance of TUNEL-positive cells (Figure 6B). Interestingly, glomerular synaptopodin levels were preserved in P13 and P28 GRP170^Pd-/-^ mice, suggesting that despite slit diaphragm disintegration, foot processes persisted in the deteriorating glomeruli. Overall, these data indicate that Grp170, by maintaining ER homeostasis, helps protect the glomerulus from proteotoxic injury. These data also suggest that defects in Grp170 function or other insults that compromise ER molecular chaperone function could contribute to diseases associated with podocyte injury.

**Figure 6.**
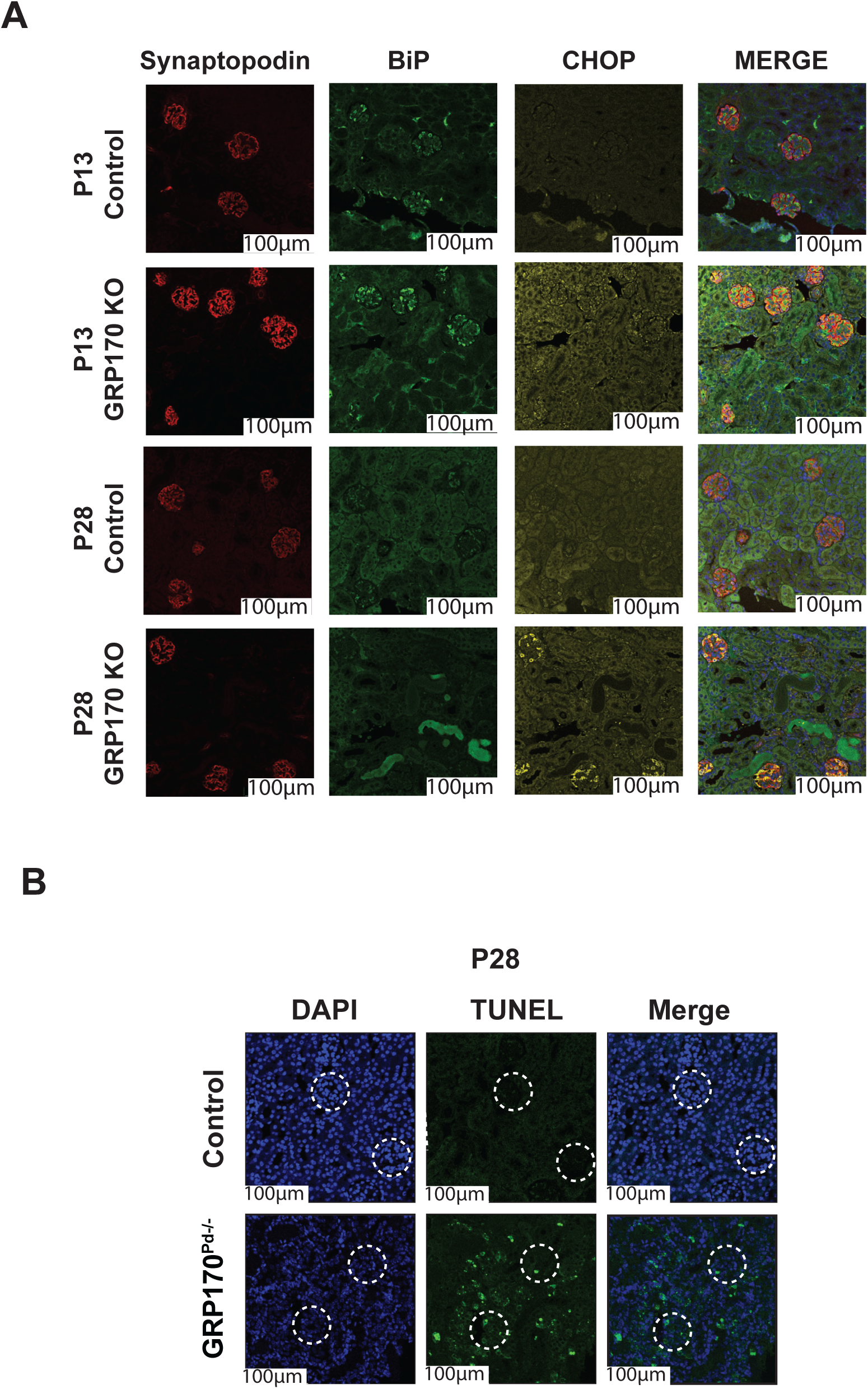
The unfolded protein response is activated in GRP170^Pd-/-^ mice. (**A**): Kidney sections from GRP170 deficient and control mice at P13 and P28 were subjected to synaptopodin (podocyte marker), BiP and CHOP staining. Representative images are shown. Scale bars: 100 μM. (**B**) TUNEL staining of P28 GRP170^Pd-/-^ and control mice. Glomeruli are identified by dashed circles. Scale bars: 100 µM.

## DISCUSSION

Molecular chaperones aid in protein folding, retain non-native proteins in a soluble state, and help deliver aberrant proteins to proteolytic pathways and are therefore indispensable to maintain cellular and organismal proteostasis(Brandvold and Morimoto 2015; Hendershot et al. 2023). Impaired chaperone function has been implicated in myriad diseases affecting nearly every organ, including the kidney (Wang and Kaufman 2016; Shukla and Narayan 2025).

Moreover, the transcription of numerous ER-localized chaperones is induced by the UPR. To date, research to understand the mechanism of chaperone function in the ER and the UPR has largely relied on pharmacologic inducers and modulators of the UPR. However, chemical UPR inducers such as tunicamycin and thapsigargin are non-specific and cause widespread toxicity. To overcome these secondary effects, we generated a genetic system with which to study how the UPR affects glomerular function and the contribution of a specific molecular chaperone, GRP170, in protecting the glomerulus from proteotoxic injury. GRP170 is ideally positioned to act as a protective chaperone due to its multi-faceted role as both a modulator of BiP, the primary Hsp70 in the ER, and as a bona fide “holdase” for misfolded proteins(Behnke et al. 2015; Behnke and Hendershot 2014; Park et al. 2003; Behnke et al. 2016). In addition, because loss of GRP170 in other model systems results in induction of the UPR, and the UPR has been linked to glomerular disease, we generated a constitutive podocyte-specific GRP170 knockout mouse (Kitano et al. 2004; Zhang et al. 2025; Porter, Nguyen, et al. 2022; Mann et al. 2024; Cybulsky 2017; Cybulsky et al. 2011). We show that GRP170 is essential for maintenance, but not development, of the glomerulus. Although the GRP170-deficient mice initially appeared healthy, chaperone loss was accompanied by progressive podocyte injury, including foot process effacement and slit diaphragm deterioration, culminating in proteinuria, hypoalbuminemia, and hyperlipidemia characteristic of nephrotic syndrome. Furthermore, we discovered that the pro-apoptotic arm of the UPR was activated by 4 weeks of age.

GRP170 plays a critical role in several pathways required to support ER function and homeostasis, including protein translocation, protein folding, UPR regulation, and the disposal of misfolded ER proteins via the ER associated degradation (ERAD) pathway(Mann et al. 2024; Behnke et al. 2015; Inoue and Tsai 2016). Based on these diverse roles, it is remarkable that the knockout mice are viable and survive for a month without intervention. Their viability suggests that GRP170 is not essential for prenatal and early postnatal podocyte development. Alternatively, survival may reflect the dual nature of the UPR: while chronic or unresolved UPR ultimately leads to cell and tissue death, transient activation allows cells to cope with stress (Cybulsky 2017; Hetz et al. 2020). In this context, GRP170 deficiency could be initially nephroprotective, or UPR redundancy may temporarily compensate for GRP170 deficiency.

Prenatal survival may be aided by the low glomerular filtration rate (GFR) *in utero*, which limits proteinuria during nephrogenesis(Iacobelli and Guignard 2021). As GFR increases after birth, however, proteinuria worsens, compounding the intrinsic toxicity of impaired proteostasis(Turin et al. 2013). While the precise mechanisms remain to be fully elucidated, the severe podocyte injury caused by GRP170 deletion may stem from the functional relationship between GRP170 and BiP. In the absence of GRP170, BiP is unable to regulate misfolded protein binding and release. Consequently, BiP becomes sequestered in insoluble protein complexes within the ER and is unavailable to bind and inhibit activation of the three UPR sensors, IRE1, ATF6, and PERK (Mann et al. 2024). This, in turn, may lead to sustained UPR induction, and culminate in apoptosis (Mann et al. 2024; Behnke et al. 2015; Inoue and Tsai 2016).

Previous studies identified elevated levels of GRP170 under conditions of glomerular injury or stress. For example, GRP170 expression was elevated in primary rat podocytes treated with the ER stress-inducer tunicamycin, and GRP170 levels increased in a dose- and time-dependent manner (Inagi et al. 2005). Similarly, in a transgenic megsin overexpression rodent model mimicking IgA nephropathy, abnormal ER protein accumulation caused marked upregulation of ER chaperones, including GRP170 (Inagi et al. 2005). Elevated serum levels of GRP170 have also been documented in the streptozotocin diabetic rat model, and patients with diabetic kidney disease and albuminuria similarly present with elevated serum GRP170 compared to healthy controls (Wang et al. 2019). In addition, in a rat model of autoimmune thyroid-induced glomerulonephritis, ER stress was associated with increased expression of BiP and GRP170, along with enhanced phosphorylation of PERK and eIF2α, a downstream PERK effector, in the glomerulus. Priming the UPR by pretreatment with the ER stress inducers thapsigargin or tunicamycin amplified BiP and GRP170 expression and ameliorated glomerular injury and proteinuria, illustrating the protective role of the UPR (Inagi et al. 2008). Similarly, exogenous induction of mild ER stress in glomerular cells can reduce proteinuria, suggesting that the adaptive/protective UPR helps mitigate glomerular injury (Cybulsky 2010).

Here, we also identify a protective role for GRP170 in the podocyte, a result that aligns with prior studies indicating that a broader cohort of ER chaperones regulate glomerular susceptibility to injury. For example, mice heterozygous for a mutation in the ER luminal Hsp70, BiP, develop glomerulosclerosis and tubulointerstitial disease (Kimura et al. 2008), which is consistent with the roles played by BiP in ER physiology (see above and (Pobre et al. 2019).

Similarly, in FSGS, membranous nephropathy, and diabetic nephropathy, ER chaperones—including GRP78 (BiP) as well ERdj3 (a BiP co-chaperone) and MANF (another BiP regulator)—are upregulated in podocytes in response to misfolded protein accumulation and cellular stress (Yuan et al. 2015; Inagi et al. 2008; Tousson-Abouelazm et al. 2020; Markan et al. 2009). The use of the nonspecific but protective chemical chaperone, 4-phenylbutyrate (4-PBA), also reduces proteinuria and glomerular ER stress in experimental models of glomerulonephritis (Tousson-Abouelazm et al. 2020).

While abundant evidence links the importance of molecular chaperone function, UPR induction, and kidney disease, establishing and characterizing the causal relationship between these events is challenging at least in part because deleting molecular chaperones or UPR components is often embryonically lethal. More pertinent, only a few podocyte tissue-specific deletion models exist (Cybulsky 2017; Inagi et al. 2014; Wu et al. 2023). For example, in a podocyte-specific *IRE1α* deletion mouse model, IRE1α deficiency blocks induction of one of the two protective branches of the UPR. The IRE1α-deficient animals eventually developed progressive albuminuria at 5 months of age, but only after 9 months did these mice develop histopathologic evidence of glomerular injury, including glomerular hypertrophy, capillary enlargement, and collagen deposition and foot process effacement (Kaufman et al. 2017). However, when the *IRE1α* deletion mice were subjected to adriamycin-induced nephrosis—a proteinuric glomerular disease model mimicking FSGS—loss of IRE1 significantly exacerbated the phenotype (Xie et al. 2021; Cybulsky et al. 2024). Adriamycin treatment also provoked rapid onset of proteinuria, glomerular and podocyte degeneration, and apoptosis. In a separate study, a podocyte-specific SEC63 knockout mouse was evaluated. SEC63 is an ER localized molecular chaperone that, like GRP170, acts as a cochaperone for BiP, but is specifically required for protein translocation(Jung and Kim 2021). Unlike the GRP170^Pd-/-^ mouse model described here, renal disease was not apparent when SEC63 was absent, although the UPR was activated. However, when the UPR was further impaired by deleting XBP1, which is the target of IRE1, Pod-SEC63/XBP1 KO mice were unable to compensate for SEC63 deficiency and developed proteinuria. After 2 months, glomerular injury was observed (Kaufman et al. 2017). Interestingly, the phenotype of our GRP170^Pd-/-^ mouse is similar to but more severe than that reported for either the podocyte-specific IRE1α- or the SEC63/XBP1-deletion mice (Kaufman et al. 2017). In combination, these studies and our work illustrate the importance of ER proteostasis and the UPR on podocyte health and resistance to stress-induced injury.

We additionally observed a striking increase in circulating triglycerides and cholesterol in GRP170^Pd-/-^ mice. These results are likely a consequences of nephrotic syndrome, in which sustained proteinuria and hypoalbuminemia drive hepatic lipoprotein synthesis and impair lipoprotein catabolism, leading to hyperlipidemia (Vaziri 2016). The expression of genes governing lipid levels is known to be altered in the setting of nephrotic syndrome. For example, the gene for ApoA-1 is induced in models of nephrotic syndrome by the transcription factor EGR-1, resulting in an increase in circulating HDL cholesterol (Zaiou et al. 1998b). Recent evidence indicates that UPR activation itself can amplify lipogenic protein expression in the kidney, resulting in enhanced lipid synthesis and renal lipid accumulation, which can compromise the glomerular filtration barrier (Zhou et al. 2025; Figueroa-Juárez et al. 2021). Hence, defective UPR signaling in P28 GRP170^Pd-/-^ podocytes may impair lipid handling, thereby directly contributing to podocyte injury independent of systemic lipoprotein metabolism.

In this work, we intentionally chose to develop an *in vivo* murine model system to maximize the clinical relevance of our findings. Podocytes have a unique ultrastructure with close associations with neighboring endothelial and mesangial cells and with unique cellular stresses relating to the flow of blood and urinary filtrate through the glomerulus. These cannot be adequately replicated by any *in vitro* platform, at least not currently. However, future studies may require the use of primary or immortalized podocytes, co-culture systems, and/or human kidney organoids to fine-tune the pathway(s) that lead from ER stress to UPR induction to podocyte injury.

In summary, our podocyte-specific GRP170 deletion mouse highlights the essential role of an ER chaperone and the UPR in podocyte health. Our model also provides a valuable tool to investigate which UPR-regulated factors govern podocyte development and maintenance, as well as to ask whether drugs that modulate the UPR or ER proteostasis prevent or ameliorate glomerular disease. While the rapidity of podocyte degeneration and the likelihood that GRP170 deletion compromises postnatal nephron development may limit its utility for the study of adult-onset podocytopathies, our model may additionally be used to probe the contribution of the UPR to congenital nephrotic diseases. Furthermore, heterozygous GRP170^Pd+/-^ mice are likely more susceptible to ER stress and predisposed to UPR dysfunction. Accordingly, in the future we will leverage this podocyte injury sensitized model to clarify which specific factors within the three arms of the UPR promote or exacerbate podocyte injury.

## MATERIALS AND METHODS

### Antibodies

Unless indicated otherwise, all antibodies and antisera used in this study are indicated in Supplemental Table 1.

### Animal maintenance and lines

Podocyte specific GRP170 KO mice (C57Bl/6 background), GRP170^Pd-/-^, were generated by crossing mice with a floxed *HYOU1* allele to *NPHS2*-Cre mice which constitutively express Cre recombinase exclusively in podocytes (Porter, Nguyen, et al. 2022; Moeller et al. 2003). All experiments conformed to NIH Guide for the Care and Use of Laboratory Animals and were approved by the University of Pittsburgh IACUC. Male and female mice were used in each experiment. Mice were housed in a temperature-controlled room with a 12-hour light-dark cycle and allowed free access to standard chow and deionized water. Mouse genotyping was carried out by Transnetyx (Memphis, TN).

### Serum chemistry

Whole blood was aspirated from the right ventricle of anesthetized mice. Blood chemistries (Na^+^, K^+^, Cl^-^, BUN, creatinine, glucose, hemoglobin, and hematocrit) were monitored using an iStat (Abbott Point of Care). The upper limit of quantification for BUN is 140 mg/dl and the lower limit of detection for creatinine is 0.2 mg/dl. For statistical validation, values above or below these thresholds were considered 140 mg/dl or 0.2 mg/dl, respectively. Serum albumin was measured using a VITROS chemistry analyzer (UT Southwestern Metabolic Phenotyping Core).

### Lipid Analysis and Apolipoprotein B Detection

Plasma triglycerides were analyzed using Fujifilm Healthcare L-type Triglyceride M microtiter kit and multi-calibrator lipids (Fisher Scientific) as directed. Plasma cholesterol levels were measured using Wako Cholesterol E Colorometric kit (Fisher Scientific). Briefly, 200 µl of cholesterol E buffer was added to 10 µl of either a blank (water), standard or plasma in a microtiter plate, incubated at 37 ℃ for 5 minutes and then absorbance was analyzed at both 600 and 700 nm.

To visualize ApoB48 and ApoB100, plasma samples were subject to SDS-PAGE (5% acrylamide gel) and western blotting. Equal protein loading was determined using Revert 700 total protein stain (LicorBio) and scanned on an Odyssey (LI-COR Biosciences, Lincoln, NE). Following removal of the total protein stain, the membrane was blocked in milk containing sodium azide for 1 hour. The blocking solution was then discarded, and the membrane was incubated with an anti-ApoB primary antibody (1:1000 dilution) overnight. The membrane was then incubated in HRP-conjugated secondary antibody. Protein bands were visualized using a Bio-Rad ChemiDoc imager and quantified using ImageJ 1.54h software (NIH).

### Histopathology and Immunostaining

Kidneys were collected and fixed in formalin for immunohistochemistry (IHC) or in 4% paraformaldehyde for indirect immunofluorescence. Following fixation, tissues were paraffin-embedded and sectioned at 4 μm (IHC) or 3 μm (IF) using standard histological procedures. H&E staining was performed by the Health Sciences Core Research Facility, Pitt Biospecimen Core at the University of Pittsburgh and imaged with a 40× objective on a Leica Aperio Scanner. For indirect immunofluorescence, slides were incubated with the indicated primary antibodies (see Supplemental Table 1) in 3% BSA overnight at 4°C. Anti-BiP antibody was described by Hendershot and colleagues(Hendershot et al. 1995). After washing with PBS, slides were incubated with corresponding fluorophore-conjugated secondary antibodies diluted 1:100 in 3% BSA overnight at 4 degrees. Where indicated, sections were then counterstained with DAPI (1 µg/ml) for 10 min and all samples were preserved in mounting media (Fluoro Gel with Dabco, Electron Microscopy Sciences). Images were acquired using a 10x objective on a Leica Stellaris 8 with identical acquisition settings for all experimental groups and processed with FijimageJ. WT1 expression was quantified by counting WT1 positive cells in each glomerulus (3-9 glomeruli per slide) and dividing the result by the cross-sectional area of the glomerulus.

### Statistics

Results are presented as mean +/- standard deviation (SD) unless otherwise noted. Graphpad Prism software (version 10.5) was used for statistical analysis. When comparing two groups, a Students’ t-test was performed. For comparison of three or more experimental groups, ANOVA with *post-hoc* Tukey multiple comparison tests were used to assess significance. In all cases, a threshold of p <0.05 was considered statistically significant.

## ACKNOWLEDGEMENTS

This work was supported by NIH grant R01 DK117126 (T.M.B.), NIH grant R35 GM131732 (J.L.B.), NIH grant K08 DK136914 (A.W.P.), UPMC Children’s Research Advisory Committee Start Up Grant (A.W.P.), NIH grant R01DK131991, Veterans Affairs Research Grant I01BX005680 (R.J.T.), and NIH grant RO1HL147818 (T.R.K). Institutional support from NIH grants U54 DK137329 and P30DK079307 is also acknowledged.

---

**Figure S1:** Mesangial and endothelial cells are present in the glomeruli of GRP170^Pd-/-^mice postnatally but degenerate by P28 as glomeruli deteriorate. Kidney sections from GRP170 deficient and control mice at postnatal days 13 and 28 were subjected to (A) PDGFRβ (mesangial cell marker) and nephrin (slit diaphragm marker) staining or (B) endomucin (endothelial cell marker) and DAPI. Representative images are shown. Scale bar: 100 µm.

