## Supplementary material for "An endoplasmic reticulum resident molecular chaperone, GRP170, prevents stress-induced glomerular injury": Figure S1

Supplemental Figure S1

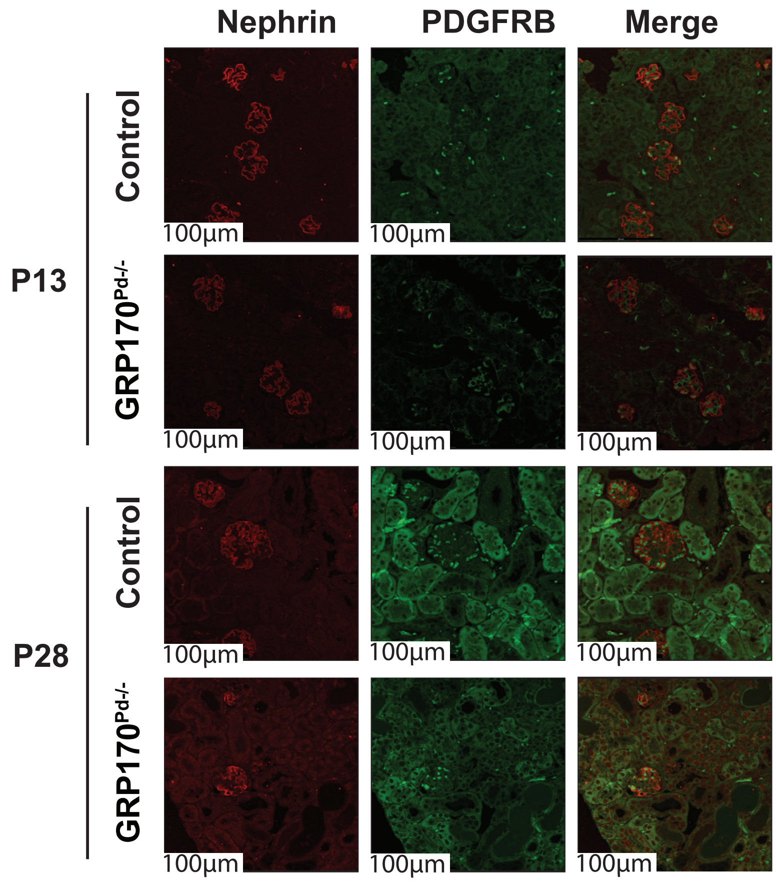


**Figure S1: Mesangial cells are present in the glomeruli of podocyte-specific GRP170 deletion mice.** Kidney sections from GRP170 deficient and control mice at postnatal days 13 and 28 were subjected to (A) PDGFRB (mesangial cell marker) and counterstained with nephrin (slit diaphram marker). Representative images are shown. Scale bar: 100 µm.
