## Supplementary material for "An endoplasmic reticulum resident molecular chaperone, GRP170, prevents stress-induced glomerular injury": Table S1

| Target | Source | Host | Species reactivity (according to manufacturer) | Application (Dilution) |
|---|---|---|---|---|
| BiP | Hendershot et al., 1995 | rabbit | N/A | IF (1:100) |
| CHOP (LC3F7) | Cell Signaling #2895S | mouse | human, mouse, rat | IF (1:200) |
| Synaptopodin (P-19) | Santa Cruz | goat | human, mouse, rat, dog | IF (1:100) |
| WT1 | Thermo fisher Scientific PA1-20991 | rabbit | rat | IF (1:100) |
| Endomucin (V.7C7) | Santa Cruz (sc-69495) | rat | mouse, rat | IF (1:100) |
| Nephrin (G-20) | Santa Cruz (sc-32530) | goat | human, mouse, rat | IF (1:50) |
| PDGFRβ (28E1) | Cell Signaling #3169 | rabbit | human, mouse, rat | IF (1:100) |
| Donkey anti-rabbit 488 | Jackson ImmunoResearch 711-545-152 | donkey | rabbit | IF (1:100) |
| Donkey anti-rat 488 | Jackson ImmunoResearch 712-547-003 | donkey | rat | IF (1:100) |
| Donkey anti-mouse 488 | Jackson ImmunoResearch 715-545-151 | donkey | mouse | IF (1:100) |
| Donkey anti-rabbit 594 | Jackson ImmunoResearch 711-585-152 | donkey | rabbit | IF (1:100) |
| Donkey anti-goat 647 | Jackson ImmunoResearch 705-605-003 | donkey | goat | IF (1:100) |
| Donkey anti-goat 594 | Jackson ImmunoResearch 705-585-003 | donkey | goat | IF (1:100) |
| ApoB (A-6) | Santa Cruz (sc-393636) | mouse | human, mouse, rabbit | WB: (1:1000) |
| Anti-mouse HRP-linked Antibody | Cell Signaling #7076 | horse | mouse | WB: (1:10000) |
